# A discovery-based proteomics approach identifies protein disulfide isomerase (PDIA1) as a biomarker of β cell stress in type 1 diabetes

**DOI:** 10.1101/2021.12.22.473924

**Authors:** Farooq Syed, Divya Singhal, Koen Raedschelders, Preethi Krishnan, Robert N. Bone, Madeline R. McLaughlin, Jennifer E. Van Eyk, Raghavendra G. Mirmira, Mei-Ling Yang, Mark J. Mamula, Huanmei Wu, Xiaowen Liu, Carmella Evans-Molina

**Author notes:** Address correspondence and requests for reprints to: Carmella Evans-Molina, MD, PhD, Indiana University School of Medicine, 635 Barnhill Drive, MS 2031A, Indianapolis, IN 46202.

## Abstract

**Background:** Activation of stress pathways intrinsic to the β cell are thought to both accelerate β cell death and increase β cell immunogenicity in type 1 diabetes (T1D). However, information on the timing and scope of these responses is lacking.

**Methods:** To identify temporal and disease-related changes in islet β cell protein expression, data independent acquisition-mass spectrometry was performed on islets collected longitudinally from NOD mice and NOD-SCID mice rendered diabetic through T cell adoptive transfer.

**Findings:** In islets collected from female NOD mice at 10, 12, and 14 weeks of age, we found a time-restricted upregulation of proteins involved in the maintenance of β cell function and stress mitigation, followed by loss of expression of protective proteins that heralded diabetes onset. Pathway analysis identified EIF2 signaling and the unfolded protein response, mTOR signaling, mitochondrial function, and oxidative phosphorylation as commonly modulated pathways in both diabetic NOD mice and NOD-SCID mice rendered acutely diabetic by adoptive transfer, highlighting this core set of pathways in T1D pathogenesis. In immunofluorescence validation studies, β cell expression of protein disulfide isomerase A1 (PDIA1) and 14-3-3b were found to be increased during disease progression in NOD islets, while PDIA1 plasma levels were increased in pre-diabetic NOD mice and in the serum of children with recent-onset T1D compared to age and sex-matched non-diabetic controls.

**Interpretation:** We identified a common and core set of modulated pathways across distinct mouse models of T1D and identified PDIA1 as a potential human biomarker of β cell stress in T1D.

## Introduction

Type 1 diabetes (T1D) results from an immune-mediated destruction of the insulin-producing β cells and manifests clinically after a threshold reduction in β cell mass and function. Data from clinical cohorts of autoantibody-positive individuals followed longitudinally suggest that β cell function is impaired very early during disease progression, often years before the onset of clinical disease (1). In parallel, histologic studies performed on pancreatic sections from organ donors with autoantibody positivity and T1D demonstrate variable reductions in β cell mass before and at diabetes onset (2–4). Findings from ex vivo disease models and pancreatic sections from human organ donors with diabetes have linked changes in β cell mass and function with the activation of a variety of β cell stress pathways that are thought to both accelerate β cell death and increase β cell immunogenicity (5–11).

At present, longitudinal imaging of the β cell compartment and sampling of the pancreas in living individuals is not clinically feasible. Studies performed using isolated islets or pancreatic sections from organ donors with diabetes provide critical information about disease pathogenesis but enable only a single snapshot view of disease pathogenesis. Improved temporal resolution of the molecular programs modulated within the pancreatic β cell during T1D evolution has the potential to inform therapeutic and biomarker strategies in humans (12–14), underscoring the need to interrogate alternative model systems.

In this regard, the non-obese diabetic (NOD) mouse has been used to study T1D pathogenesis for over three decades (15–17). Islets in NOD mice show evidence of immune cell infiltration as early as four weeks of age (18), and approximately 60-80% of female NOD mice develop diabetes by 16-20 weeks of age (15, 19). Consistent with patterns observed in humans, β cell function and mass decline during the pre-diabetes phase. In cross-sectional analyses, subsets of overlapping stress pathways have been identified in β cells from NOD mice and in human islets from organ donors with diabetes (16,20,21). To gain additional insight into the time course of molecular changes in the β cell during T1D progression, we used a data independent acquisition-mass spectrometry (DIA-MS) based approach to monitor longitudinal changes in the islet cell proteome during early and late disease progression in NOD mice. In parallel, we compared proteomic signatures in islets from diabetic NOD mice with those observed in islets from NOD-SCID mice rendered acutely diabetic by the adoptive transfer of T cells from NOD-BDC2.5 mice. Finally, to gain insight into potentially protective pathways, proteomes generated from NOD mouse islets at the time of diabetes onset were compared with those from NOD mice that remained diabetes-free through 46-48 weeks of age. Key findings were validated using immunofluorescence in pancreatic tissue sections. To illustrate the utility of this approach in prioritizing β cell proteins as T1D biomarkers, we focused on protein disulfide isomerase A1 (PDIA1) as an example of a secreted protein that was found to be differentially expressed in NOD islets during diabetes progression. PDIA1 was selected as a target for the development of a high-sensitivity electrochemiluminescence assay using MesoScale Discovery technology. Using this assay, we demonstrated increased β cell expression and circulating concentrations of PDIA1 in plasma from NOD mice during the evolution of T1D and in the serum of children with recent-onset T1D compared to age and sex-matched pediatric controls.

## Results

### Analysis of temporal changes in the NOD proteome during disease progression

To characterize temporal changes in islet protein expression during diabetes progression, pancreatic islets were isolated from age-matched CD1 and NOD mice at 10, 12, and 14 weeks of age and at the time of diabetes onset (mean age of diabetes development 17 ± 3.3 weeks; mean ± S.D.) and analyzed using LC MS/MS (**Figure 1**). An average of 1160 proteins and an average of 897 overlapping proteins were quantified in NOD and CD1 mouse islets (**Supplementary Figure 1**). Since CD1 mice are not diabetes-prone and exhibit tightly regulated blood glucose levels, we used sex and age-matched CD1 mice to normalize protein abundance in NOD islets. To identify differentially expressed proteins at each time point, results were analyzed using median normalization and a filtering criterion of a 1.5-fold change in protein abundance.

**Figure 1:**
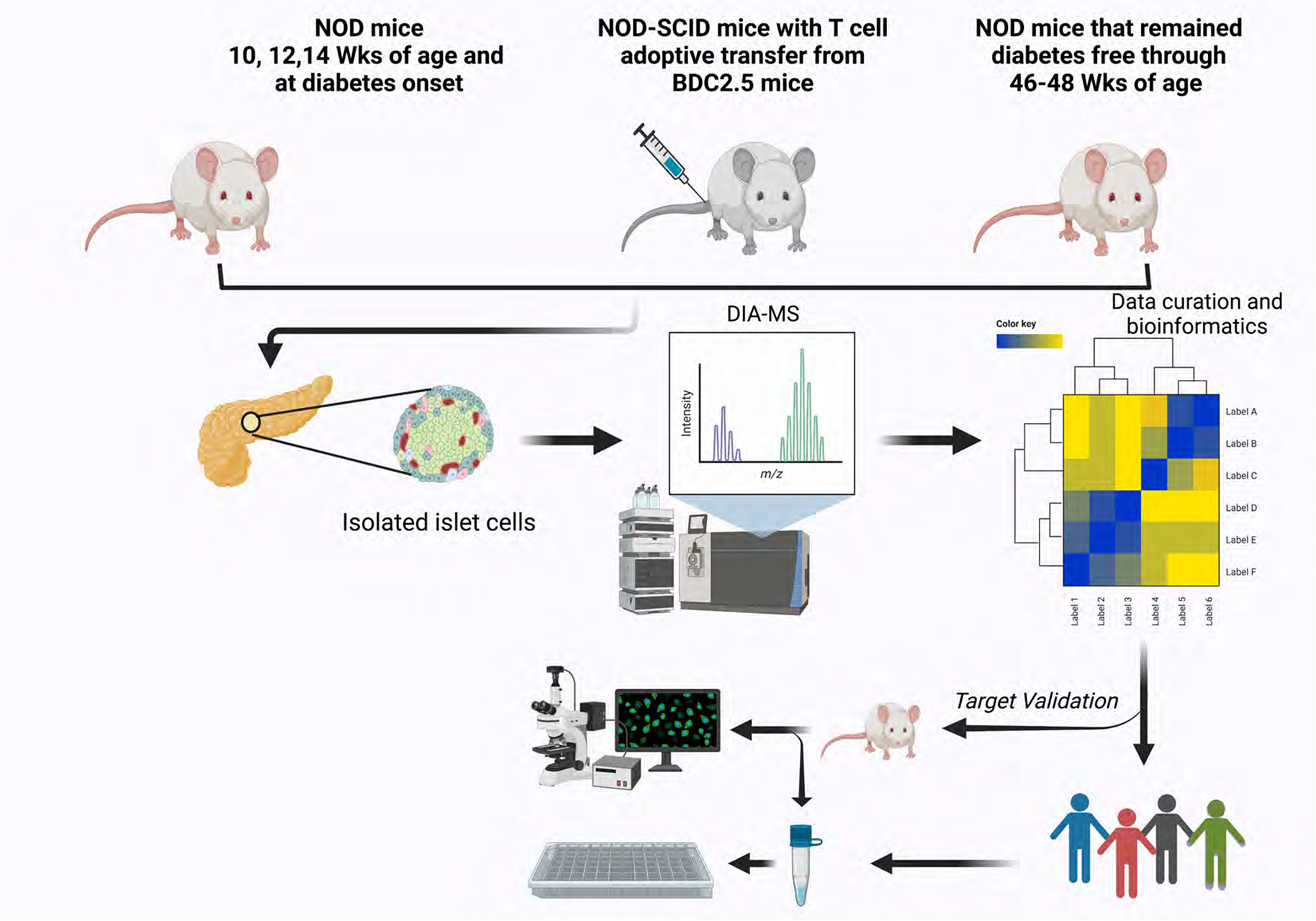
Schematic representation of study design and experimental workflow (created using BioRender.com)

In principal component analysis (PCA), NOD mice at different pre-diabetic ages (10, 12, and 14 weeks) clustered primarily as one group, whereas diabetic mice were distinctly clustered (**Figure 2A**). Unsupervised hierarchical clustering analysis of the top 30 differentially expressed (DE) islet proteins revealed notable changes in the temporal expression of several proteins during diabetes progression (**Figure 2B**). As expected, there was a time-dependent loss of insulin and islet amyloid polypeptide (IAPP) expression. Interestingly, we observed a time-restricted upregulation of several proteins implicated in stress mediation and the maintenance of normal β cell function during the prediabetic phase. Proteins exhibiting this pattern of expression included actin-related protein 2/3 complex 2 (ARPC2), which regulates actin cytoskeleton-mediated transport of secretory vesicles (22–24), collagen 1A1 (COL1A1), and collagen 1A2 (COL1A2), which are extracellular matrix proteins (25, 26), and the metallothioneins MT1 and MT2, which have been linked with suppression of immune responses (27, 28). A similar pattern was observed for peroxiredoxin 3 (PRDX3), a protein that has been linked with the regulation of mitochondrial function, and 14-3-3b, which plays a role in a number of metabolic processes, including mTOR signaling, amino acid metabolism, and mitochondrial function (29, 30). Protein disulfide isomerase A1 (PDIA1) was upregulated similarly during weeks 12 to 14. Notably, PDIA1 is a thiol reductase that plays a critical role in proinsulin folding and regulation of ER function (31). Overall, this pattern of upregulation was observed through the 14-week timepoint followed by a declining pattern expression of several of these proteins with potentially protective roles that heralded diabetes development (**Figure 2B**).

**Figure 2:**
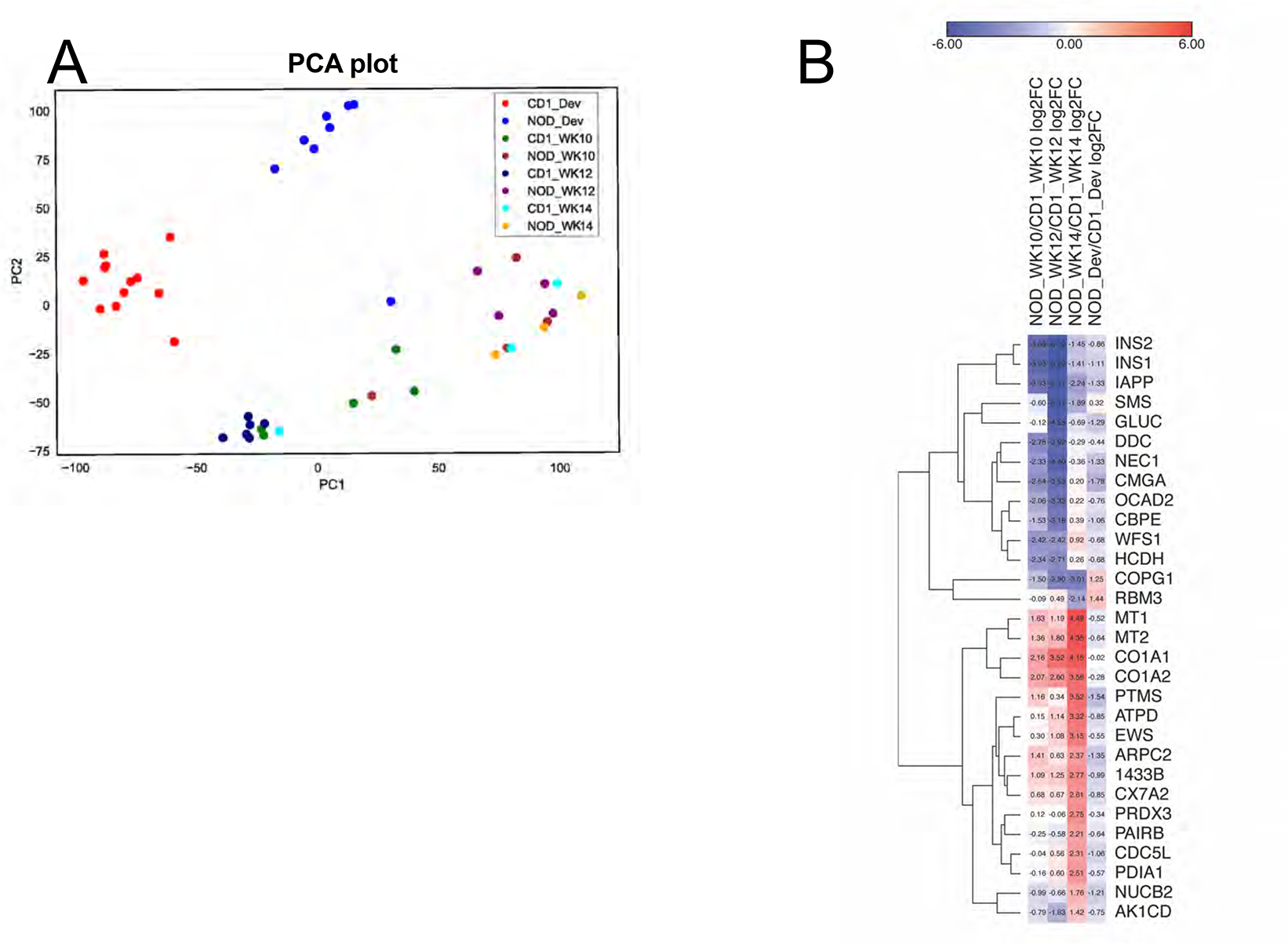
Proteomic analysis of pancreatic islets in NOD and CD1 mice over time. (A) Principal component analysis (PCA) of all quantified islet proteins from NOD and CD1 mice at 10, 12, and 14 weeks of age and during diabetes onset indicates the reproducibility of biological replicates. (B) Unsupervised hierarchical clustering analysis of the top 30 differentially expressed proteins in NOD when compared to age and sex-matched CD1 mice from 10, 12, and 14 weeks of age, and during diabetes onset. (n= 3-12 animals/ group)

**Figure 3A** shows the top 10 upregulated pathways, while **Figure 3B** shows the top 10 downregulated pathways in longitudinal analysis of islets from NOD mice using ingenuity pathway analysis. During diabetes progression, pathways related to Cdc42, integrin signaling, actin, epithelial adherens, and mTOR signaling were all upregulated (**Figure 3A**). EIF2 signaling, which is involved in global mRNA translation initiation and is a target during the unfolded protein response and ER stress (32, 33), was markedly upregulated at weeks 12 and 14 and in diabetic mice (**Figure 3A**), and changes in mitochondrial function were represented in both up-and down-regulated pathways. Significantly downregulated pathways encompassed several metabolic pathways, including the TCA cycle, oxidative phosphorylation, fatty acid oxidation, and glutathione redox reactions. Sirtuin signaling and phagosome maturation were also downregulated (**Figure 3B** and **Supplementary Figures 2 and 3**).

**Figure 3:**
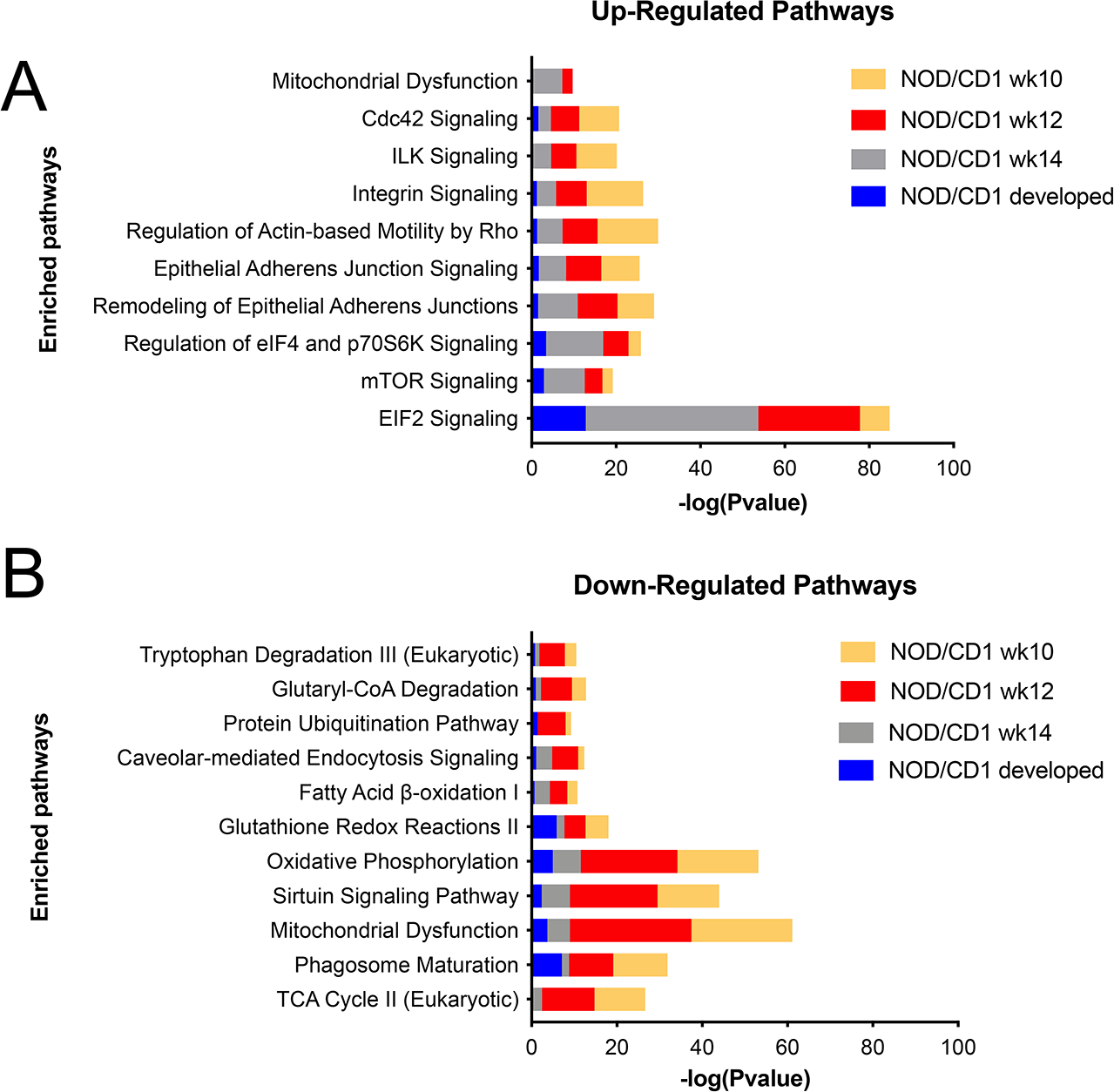
Ingenuity pathway analysis of islet proteome. The top 10 upregulated (A) and downregulated (B) canonical pathways modulated during diabetes progression in islets collected from NOD mice compared to age-matched CD1 mice.

### Comparison of spontaneous and induced models of immune cell infiltration

To identify commonalities and differences in the islet proteome between the chronic, spontaneous NOD model and an aggressive, acute model of immune-mediated β cell destruction, we compared proteomics results from islets isolated from diabetic NOD mice and islets isolated at the time of diabetes development from NOD-SCID mice that had undergone adoptive transfer of CD4+ T-cells from NOD.BDC2.5 mice. Mice in the latter group develop significant hyperglycemia around 7 days following adoptive transfer. Despite this difference in the time-course of diabetes development compared to the chronic and slowly progressive NOD model, approximately ∼65% of identified proteins were common to both models (**Figure 4A**). In addition, a comparison of functional canonical pathways suggested that similar pathways were activated in both models. Key overlapping pathways included modulation of EIF2 signaling and the unfolded protein response, mitochondrial dysfunction, oxidative phosphorylation, and mTOR signaling (**Figure 4B**). These results suggest that irrespective of the type of inflammation (acute or chronic), similar patterns of β cell stress are activated, underscoring the importance of this core set of pathways in T1D pathogenesis.

**Figure 4:**
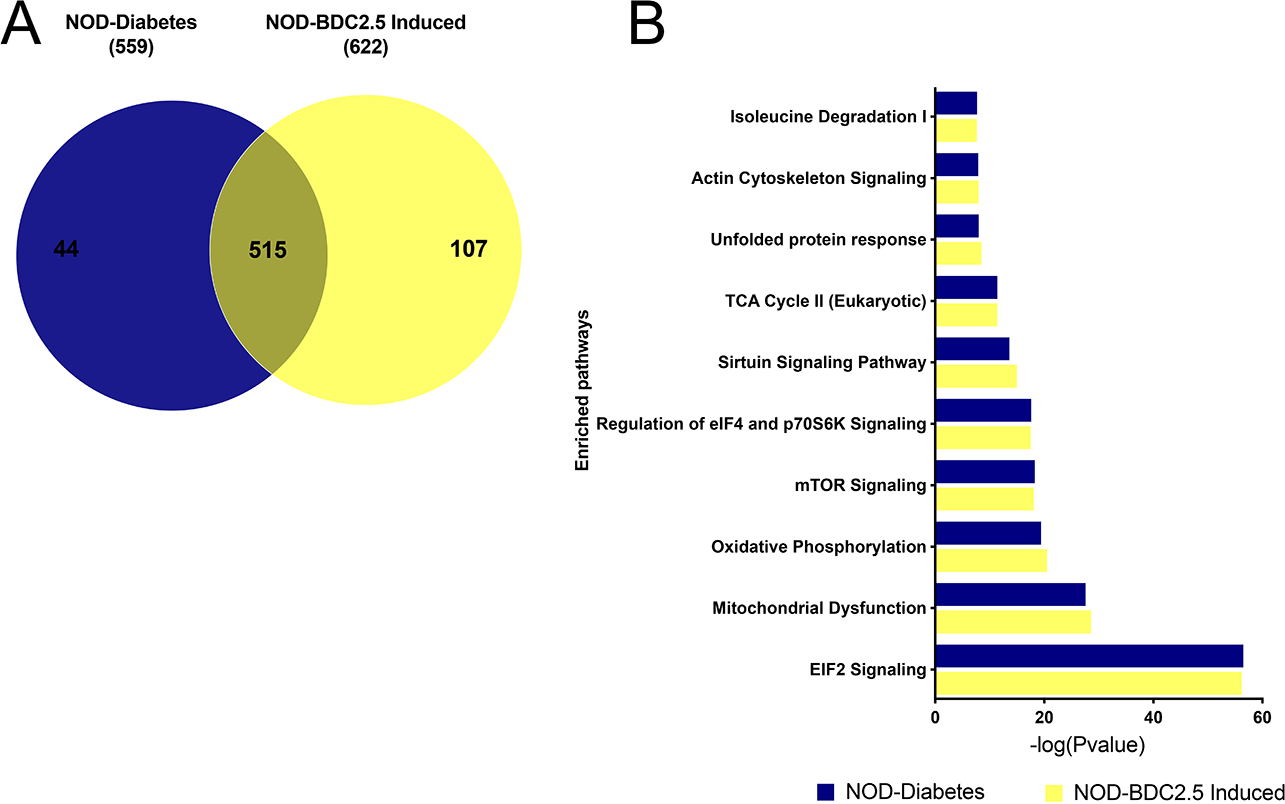
Comparison of the islet proteome between acute and chronic models of T1D. (A) Venn diagram showing protein overlap between the spontaneous and inducible models of T1D. (B) Pathway enrichment analysis identified common pathways that are significantly activated during the pathogenesis of T1D.

### Proteome comparison of NOD mice that developed diabetes and that remaining diabetes-free

We reasoned that comparing diabetic NOD mice and NOD mice that remained diabetes-free through extended follow-up might highlight protective pathways within the β cell during immune activation. Therefore, proteomic analysis was performed on islets from 46-48 wk old NOD mice who remained diabetes-free, and results were compared to islets collected from NOD mice at the time of diabetes development. Principal component analysis indicated a clear separation between the diabetes-resistant group and diabetic NOD mice (**Figure 5A**). Next, unsupervised hierarchical clustering analysis was performed using the Euclidian distance and average linkage method (**Figure 5B**). Data from this analysis revealed upregulation of several unique proteins previously linked with the mitigation of β cell stress and maintenance of normal β cell function in the diabetes-resistant NOD mice. Among the top proteins upregulated in diabetes resistant mice and down-regulated in diabetic mice were IAPP and antioxidant-1 (ATOX1), a copper chaperone shown to be protective against hydrogen peroxide and superoxide mediated-oxidative stress (34). Other key proteins showing this pattern of expression were proteasome subunit beta 10 (PSB10), which is involved in the maintenance of protein homeostasis (35), coactosin like protein (COTL1), an F-actin-binding protein that plays a role in cellular growth (36); and S100A4, which functions as an intracellular cytosolic calcium sensor (37, 38).

**Figure 5:**
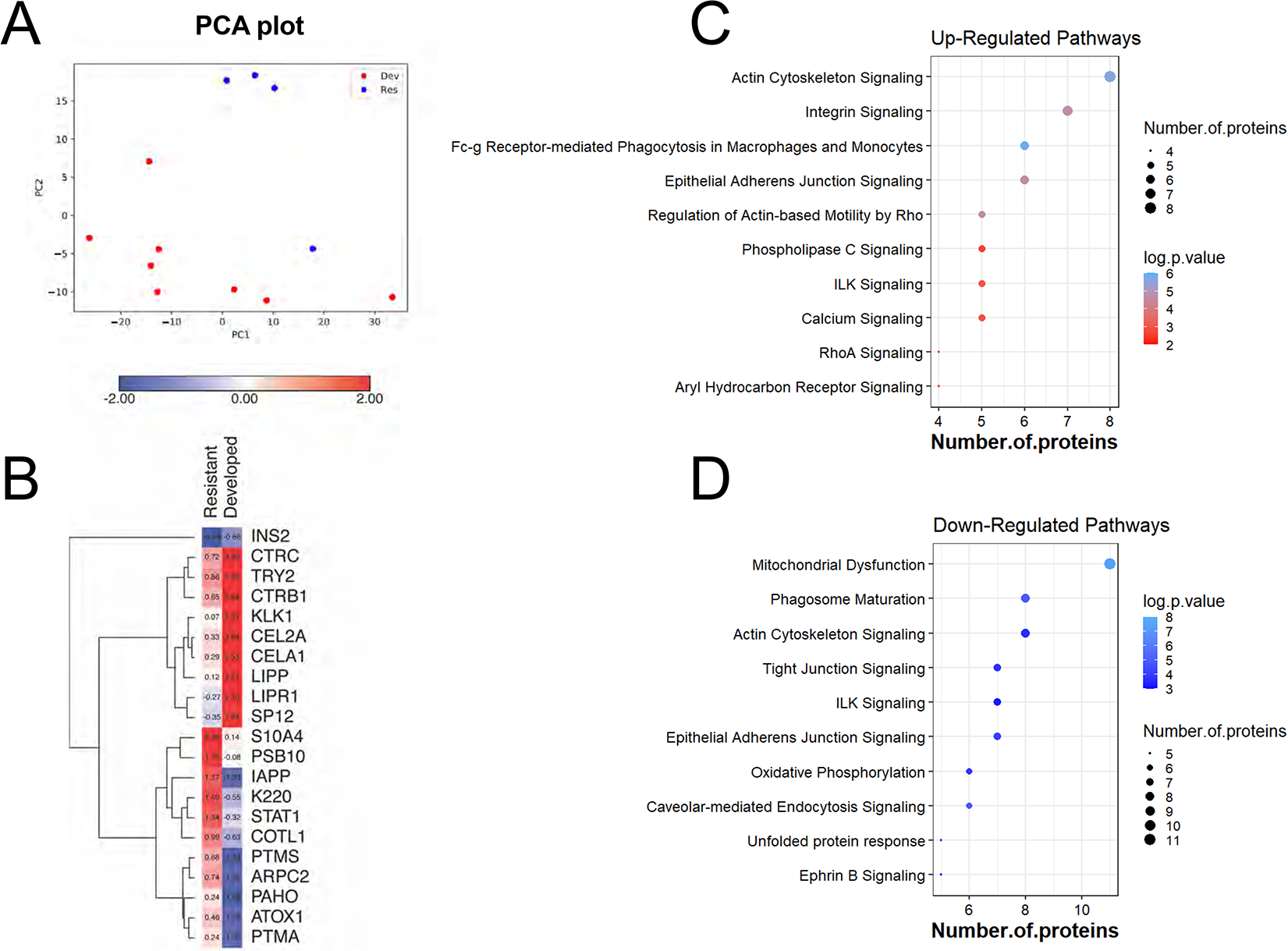
Proteomics analysis of diabetes-resistant NOD mice and NOD mice that developed diabetes. (A) Principal component analysis (PCA) of the islet proteome in NOD mice that remained diabetes-free through 46-48 wks of age (Res) and NOD mice at the time of the development of diabetes (Dev). (B) Heatmap showing top differentially expressed islet proteins in diabetes-resistant mice and mice at the time of diabetes onset. Shown are up-regulated (C), and down-regulated (D) pathways (y-axis) and the corresponding number of proteins (x-axis) differentially expressed in islets from diabetes- resistant mice.

Pathway enrichment analysis was performed to uncover the signaling pathways that were differentially regulated between these two groups. **Figure 5C** shows the top 10 significantly upregulated pathways in diabetes-resistant NOD mice compared to diabetic NOD mice. The size of each circle indicates the number of proteins enriched in each pathway, and the density of each circle represents their p-values. In diabetes-resistant NOD mice, there was a notable modulation of pathways involved in maintaining cellular homeostasis, tissue repair, tissue clearance (i.e., phagocytosis in macrophages and monocytes), and aryl hydrocarbon receptor signaling, which has been linked with the mitigation of insulitis in NOD mice. (39). In line with this, we observed downregulation of pathways related to mitochondrial dysfunction, phagosome maturation, oxidative phosphorylation, and the unfolded protein response (**Figure 5D**). Interestingly, actin cytoskeleton signaling and epithelial adherens junction signaling were identified as being among both the up-and down-regulated pathways, with distinct proteins implicated within the up-and down-regulated categories.

### Validation of protein targets by immunofluorescence and electro-chemiluminescence

To validate key findings from the proteomic analysis, immunofluorescence staining of pancreatic tissue sections was performed using a separate cohort of NOD mice aged 9 to 13 weeks and at diabetes onset. Three protein targets, PDIA1, 14-3-3b, and PRDX3, were selected for validation experiments based on top hits from the analysis shown in **Figure 2B** and their known roles in maintaining β cell function (30,31,40,41). Consistent with proteomics data, the staining intensity of islet PDIA1 (**Figure 6A**) and 14-3-3b increased from 9-13 weeks in NOD mice, followed by a significant loss of target protein expression at T1D onset (**Figure 6B**). Changes in the staining intensity of PRDX3 did not reach change significantly during the prediabetic timepoints (**Figure 6C**). However, we observed a significant decrease in PRDX3 at T1D onset in NOD mice.

**Figure 6:**
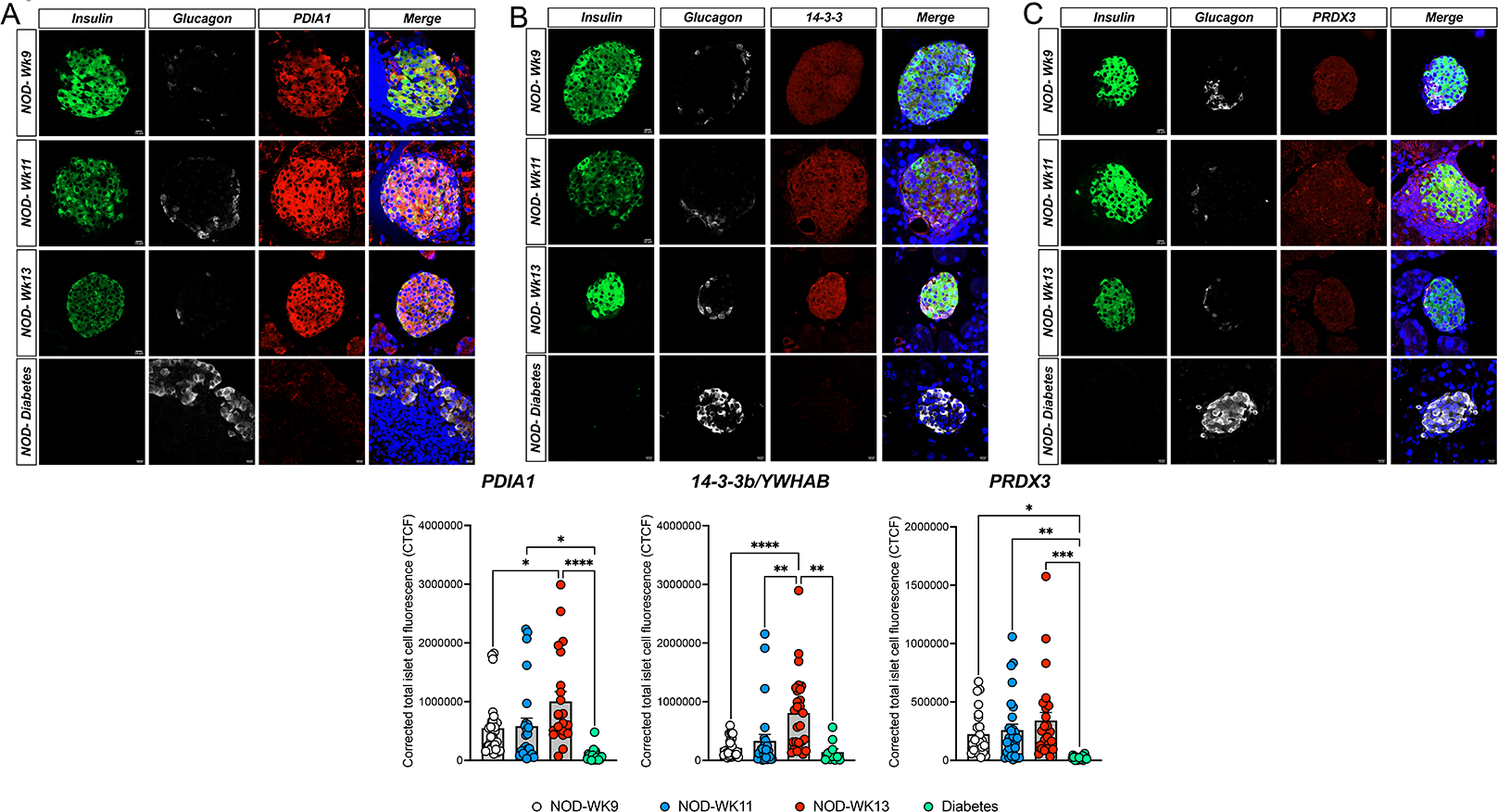
Immunofluorescence-based validation of protein targets in NOD mice. (A-C) Pancreas sections from 9, 11, and 13 wk old NOD mice and in NOD mice that developed diabetes were immunostained for PDIA1 (A), 14-3-3b (B), and PRDX3 (C) (red), and costained with insulin (green) and glucagon (white). Scale bar = 10 um. The bar graphs show the quantitation of fluorescence intensity for each protein target calculated using the corrected total islet cell fluorescence. (N=4-5 mice/age-group; 5-10 islets/mice were used for quantification; *P<0.05, **P<0.001)

### Analysis of circulating PDIA1 as a T1D associated biomarker

In addition to its intracellular role as a thiol reductase (31,42,43), PDIA1 is known to be a secreted protein (31). At present, circulating biomarkers that reflect the health of the β cell are lacking. Therefore, to determine whether the islet-specific upregulation of PDIA1 identified in the proteomics and immunofluorescence analyses was linked with changes in circulating PDIA1 levels and to test its utility as a biomarker of β cell stress in T1D, we developed a high-sensitivity electrochemiluminescence assay using Meso Scale Discovery technology. PDIA1 was measured using serially diluted (1:4) recombinant PDIA1, and this analysis showed that PDIA1 could be detected in the range of 0.152 ng/ml up to 2500 ng/ml (**Figure 7A**). Using plasma collected from the same mice employed in the longitudinal proteomics analysis shown in **Figure 2**, we found that the plasma levels of PDIA1 were significantly increased in pre-diabetic NOD mice compared to CD1 mice at 10 and 14 weeks of age (**Figure 7B-D**). However, PDIA1 levels were not different between NOD mice at the time of diabetes onset and age-matched CD1 mice (**Figure 7E**). In addition, PDIA1 levels were below the detectable range in plasma from diabetic NOD-SCID mice that had undergone T-cell adoptive transfer.

**Figure 7:**
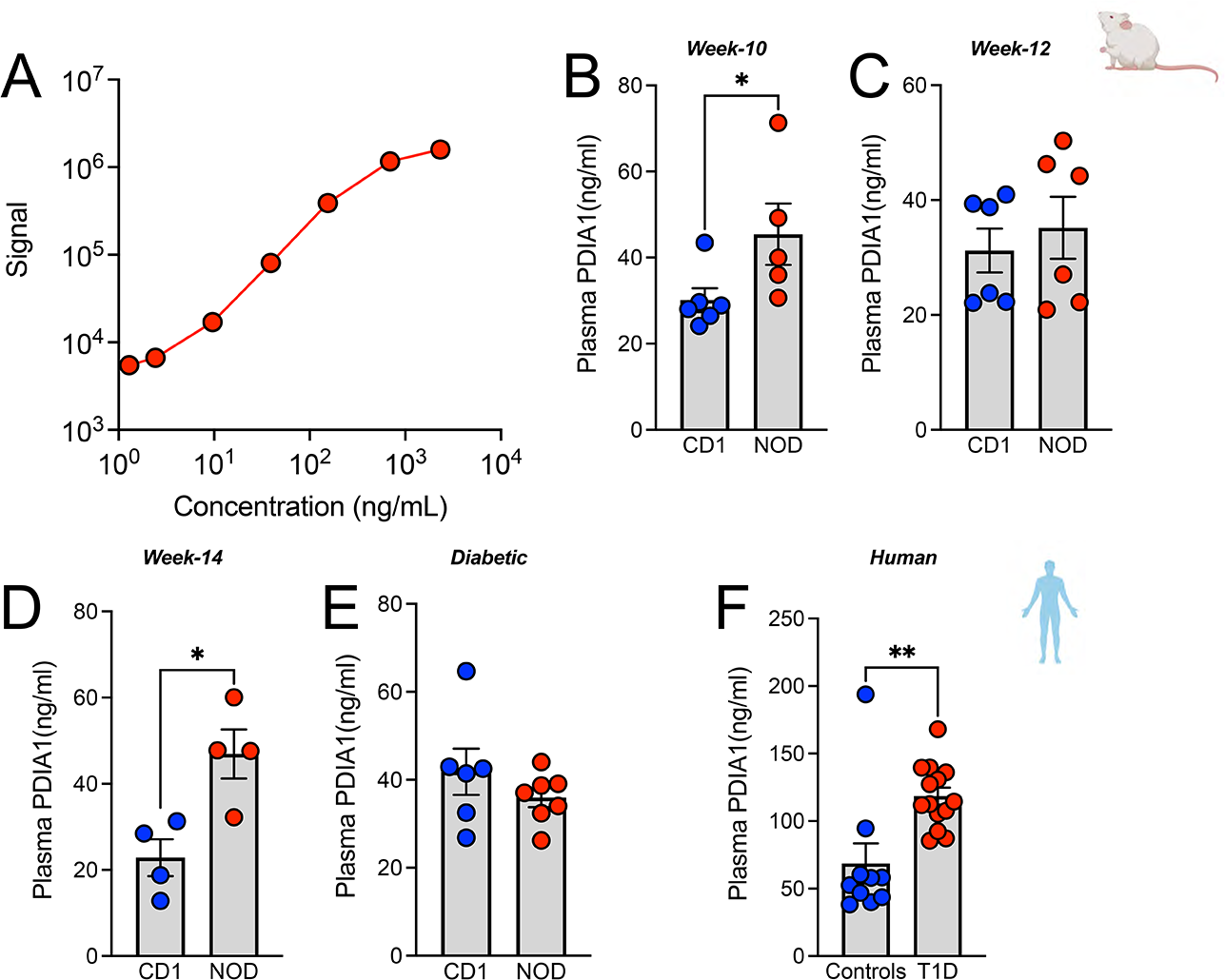
PDIA is increased in pre-diabetic NOD mice and in recent-onset T1D. (A) Standard curve generated using a serial dilution of recombinant PDIA1 protein showing higher and lower detection limits of assay sensitivity. (B-E) Measurement of circulating PDIA1 in plasma samples of sex and age-matched CD1 and NOD mice at different ages. N=4-7 mice/group, *P<0.05, **P<0.001. (F) Plasma concentration of PDIA1 in non-diabetic control subjects (n= 10, average age 12 ± 4.2 yrs) and pediatric subjects with recent-onset T1D (n=14, average age 11.57 ± 4.05 yrs) **P=0.001.

Next, we applied this assay to serum samples collected from children within 48 hours of the clinical onset of T1D (n=10; average age (mean ± SD) = 11.57 ± 4.05 yrs; 6 male; 4 female) and in serum collected from non-diabetic pediatric controls (n=13; average age = 12.1 ± 4.20; 7 male; 6 female). Interestingly, serum levels of PDIA1 were significantly higher in pediatric subjects with recent-onset T1D compared to controls, suggesting PDIA1 may have utility as a clinical, human T1D biomarker. (**Figure 7F**).

## Discussion

In this study, we aimed to identify temporal changes in islet β cell protein expression during the evolution of T1D using three distinct mouse models of T1D and high-throughput DIA-MS-based proteomics. We analyzed the proteome of islets collected from female NOD mice at three pre-diabetic time points and the time of diabetes onset and compared protein abundance with sex- and age-matched non-diabetic CD1 mice. To compare chronic and acute models of T1D, we analyzed the proteome of islets from diabetic NOD mice and NOD-SCID mice that has been rendered acutely diabetic following the adoptive transfer of T cells from NOD.BDC2.5 mice. To gain insight into potential mechanisms that contribute to β cell resiliency in the face of immune activation, we compared the islet proteomes of NOD mice at diabetes onset to NOD mice that remained diabetes-free after 46-48 weeks of observation. Finally, to illustrate the utility of this approach in prioritizing β cell proteins as T1D biomarkers in humans, we focused on protein disulfide isomerase A1 (PDIA1) as one example of a secreted protein that was differentially expressed in NOD islets during diabetes progression for the development of a high-sensitivity electrochemiluminescence assay to be employed in serum and plasma. Proteomics analysis of the three mouse models revealed several notable themes.

In the dataset obtained from the longitudinal NOD cohort, we observed an early but time-restricted increase in the expression of several proteins previously linked with secretory function, proinsulin folding, and stress mitigation, including proteins known to be involved in the mitigation of endoplasmic reticulum and oxidative stress. Interestingly, we observed week 14 (44) as a potential inflection point, where the loss of expression of these protective proteins heralded T1D onset. Consistent with this, canonical pathway analysis of differentially expressed proteins from wks 10, 12, and 14 and diabetes onset identified upregulation of pathways associated with defective insulin synthesis and several β cell stress pathways, including mitochondrial dysfunction, ER stress, and UPR activation. We observed also a downregulation of signaling pathways that were crucial for the mitigation of ongoing cellular stress, including glutathione redox signaling, which is known to counteract the effects of reactive oxygen species(45, 46) and phagosome maturation, which is involved in the clearing of cellular debris (47, 48).

This biphasic pattern is reminiscent of metabolic data from natural history cohorts of autoantibody-positive individuals who progress to T1D, where there are compensatory changes in the architecture of insulin secretion that largely maintain glycemia until ∼12 months prior to disease onset, followed by marked loss of insulin secretion and rapidly worsening glycemic control until diabetes diagnosis (1). Our findings are also consistent with cross-sectional studies that have analyzed gene and protein expression patterns in pancreatic sections from human donors with diabetes (49–52) and in previous studies in mouse models of diabetes, where a prominent role for ER and mitochondrial dysfunction has been identified during disease progression (6,10,53,54). Notably, we found these pathways are activated early in the disease process, and there is continued overlap between several of these key activated stress pathways in the NOD mouse model and in the acute, inducible model of T1D at the time of diabetes onset. While the former is a spontaneous model and the latter is an acute model of islet destruction, similarities between the proteomic analysis of these two models highlight the importance of this core set of pathways in T1D pathogenesis.

To validate selected findings from the proteomics analysis, we performed immunofluorescence experiments, focusing on three targets identified in the longitudinal NOD cohort: 14-3-3β, PRDX3, and PDIA1. Members of the 14-3-3 protein family have been implicated in various metabolic signaling pathways and have been linked with protection against apoptosis in pancreatic β cells (30). PRDX3 prevents mitochondrial dysfunction, and its overexpression has been shown to be protective against oxidative stress induced by insulin resistance and hyperglycemia (40, 55). PDIA1 is a highly abundant ER localized thiol oxidoreductase that has been implicated in glucose-stimulated insulin secretion, proinsulin processing, and protection against ER stress (31). Of note, PDIA1 has been described as a secreted protein, and in other cell types, PDIA1 release is increased in the setting of injury and stress (56, 57). Extracellular PDIA1 has been linked with the regulation of thrombus formation during vascular inflammation (58, 59), but a complete understanding of the extracellular role of this protein is lacking. Interestingly, anti-PDIA1 antibodies have been identified in patients with recent-onset T1D (60), suggesting that β cell derived PDIA1 serves as a T1D autoantigen. Against this background, we hypothesized that increased β cell expression of PDIA1 may be reflected in the circulation and that measurement of PDIA1 may have utility as a T1D biomarker. To test this possibility, we developed a high sensitivity electrochemiluminescence assay to measure serum and plasma PDIA1. Using this assay, we documented an increase in plasma PDIA1 in pre-diabetic NOD mice and in the serum of children with recent-onset T1D. To our knowledge, this represents the first assessment of circulating PDIA1 in individuals with diabetes.

Our study has several limitations that should be acknowledged. First, there are significant differences between mouse models of T1D and human T1D (61–63), underscoring the need to cross-validate findings from mouse models in human samples. We followed such a workflow and strategy with PDIA1 and documented increased serum levels of PDIA1 in a small cohort of pediatric subjects with recent-onset T1D. While biomarkers with the ability to non-invasively monitor β cell stress are notably lacking in T1D, it is important to acknowledge that ours is a small cross-sectional study and validation in larger cohorts should be performed. Whether PDIA1 is purely a marker of β cell stress or may reflect β cell mass changes is not clear from our study and should be tested in follow-up. Along these lines, it will be essential to test PDIA1 levels in samples collected from clinical cohorts followed longitudinally during T1D progression. Such an analysis will provide necessary insight into whether PDIA1 can predict T1D risk. It is interesting that PDIA1 levels were higher in children with new onset T1D, whereas plasma elevations in PDIA1 were most notable at pre-diabetic timepoints in NOD mice. This is consistent with more recent data showing that there is substantial β cell mass remaining in humans at T1D onset ((64, 65), whereas in both of the mouse models studied here, β cells are nearly completely destroyed by the time of diabetes onset (66, 67)..

Notwithstanding these limitations, our study highlights the value of unbiased proteomics approaches for identifying key β cell pathways involved in the temporal evolution of T1D. Utilizing this strategy, we identified a common set of modulated pathways across several distinct mouse models of T1D and identified PDIA1 as a potential T1D associated biomarker.

## Materials and Methods

### Animals and experimental procedures

Female NOD/ShiLTJ (NOD), NOD-BDC2.5 and NOD-SCID mice were purchased from the Jackson Laboratory. Female outbred CD1 mice were purchased from Charles River Laboratories. Mice were maintained at the Indiana University School of Medicine Laboratory Animal Resource Center under pathogen-free conditions and protocols approved by the Indiana University Institutional Animal Care and Use Committee.

Mice were allowed to be acclimated for at least two weeks upon arrival and before the initiation of experiments. Pancreatic islets were isolated at the indicated time points, or the pancreas was harvested at the indicated time points, as described previously (68, 69). Islets were hand-picked, washed twice with PBS, and stored as pellets at −80°C until use. Blood destined for plasma analysis was obtained at the time of euthanasia via cardiac puncture, transferred to a Becton Dickinson Vacutainer K2EDTA tube (Cat# 365974), and centrifuged at 5,000 rpm for 10 minutes at 4°C. The separated plasma samples were aliquoted into 1.5ml cryotubes and stored at -80°C until use.

Single-cell splenocyte suspensions were prepared for adoptive transfer experiments from 12-week-old male NOD-BDC2.5 mice, as previously described (70). CD4+ T cells were purified by negative selection (Cat# 558131, BD Biosciences), activated in 6-well plates (5×10^6^ cells/well), coated with anti-CD3 and anti-CD28 antibody (1 mg/mL each), and expanded for 72 h in T-75 flasks (5×10^6^ cells/flask) in complete RPMI 1640 medium (1% penicillin/streptomycin and 10% FBS) containing 100 U/mL IL-2. Cells were subsequently collected, washed twice with Hanks’ balanced salt solution (HBSS), and diluted to 5×10^6^ cells/mL in HBSS (71). Recipient 8-week-old immunodeficient male NOD-SCID mice received 1×10^6^ T cells via intraperitoneal injection, and blood glucose was measured daily for 21 days. Age-matched NOD-SCID mice that received HBSS alone were used as controls. The onset of diabetes was defined as two consecutive blood glucose readings of ≥ 275 mg/dL.

### Mass Spectrometry sample processing

Islet pellets were lysed and denatured by adding 48 mg of urea to ∼100 μL of pelleted cells. Lysates were ultrasonicated with 5 successive 10s pulses to ensure complete lysis and to shear DNA. Protein content was measured by BCA assay, (Pierce) and 50 μg of protein was transferred to a 1.5-mL tube and the volume was adjusted to 250 μL using 50 mM ammonium bicarbonate (pH 8.0). Sample were subsequently reduced with 25 mM of freshly prepared tris(2-carboxyethyl) phosphine at 37°C for 40 min, alkylated with 10 mM of freshly prepared iodoacetamide for 40 min at room temperature in the dark, and diluted to 800 μL with 50 mM ammonium bicarbonate. The pH of the sample was adjusted to 8.0, and digested using a 50:1 ratio of protein:trypsin (Sequence grade, Promega) at 37°C overnight in the presence of 10% acetonitrile with constant agitation, using trypsin at a 50:1 ratio. The digest was then acidified with 10% Formic Acid (pH 2-3), desalted on a 96-well microelution plate (Oasis HLA, Waters), and dried before mass spectrometry (MS) analysis.

### Data-independent acquisition LC-MS/MS analysis

Discovery proteomics was performed by liquid chromatography-tandem mass spectrometry (LC-MS/MS) using an Eksigent 415 HPLC system equipped with an Ekspert nanoLC 400 autosamplers coupled to a SCIEX 6600 TripleTOF mass spectrometer (SCIEX, Framingham, MA) operated in data-independent acquisition (DIA) mode. Two microgram of peptides were injected onto a ChipLC trap-elute system equipped with a 15-cm, 75-μm inner-diameter C18 column (300 Å stationary phase) and separated at a flow rate of 500 nL/min using a linear AB gradient of 3-35% solvent B (0.1% FA in acetonitrile) for 60 min, 35-85% B for 2 min, hold at 85% B for 5 min, and re-equilibration at 3% B for 7 min. Mass spectra were obtained with 64 variable-width precursor isolation windows. Dwell-times in MS1 and MS/MS were 250 and 45 ms, respectively, for a total cycle time of 3.2 s. The collision energy was optimized for an ion *m/z* centered on the isolation window, with the collision energy spread ranging from 5-15. Source gas 1 was set to 3, gas 2 was set to 0, curtain gas was set to 25, source temperature was set to 100°C, and source voltage was set to 2400 V.

### Post-acquisition analysis

#### Peptide library generation

Individually acquired DIA files were processed using the Signal Extraction module of the DIA-Umpire software tool (DIAu-SE). A protein sequence database was built by concatenating the target SwissProt mouse proteome database (canonical sequences), Biognosys iRT peptides for retention time alignment (Biognosys, Schlieren, Switzerland), and a random decoy sequence database with the same size as the target database for false discovery rate (FDR) estimation. Pseudospectra generated in the DIAu-SE step were then searched against the concatenated database for peptide identification. Identified peptides filtered with a 1% peptide-level FDR were used for library generation.

#### Quantitation of individual specimen DIA-MS files

Raw intensity data for peptide fragments were extracted from DIA files using the open-source openSWATH workflow against the sample-specific peptide spectral library described above. Briefly, peptide assay peak groups were extracted from raw DIA files and scored against the peptide- and decoy-spectral libraries with the same size based on a composite of 11 data-quality subscores. Target peptides with a 1% FDR were included for downstream analyses.

#### Curating files for quality

All files were individually curated prior to protein-level roll-up and subsequent quantitation. The following parameters were considered: total ion chromatogram profile and intensity, file quality within the library build (Q1, Q2, Q3 data distribution from DIAumpire), and raw distribution of proteins compared to decoys derived in openSWATH. Files exhibiting aberrant or low-quality results for any of these parameters were excluded from subsequent analysis steps. All steps were performed while blinded to filenames or experimental group.

#### Data normalization, protein-level roll-up, and statistical analyses

The total ion current associated with the MS2 signal across the chromatogram was calculated for normalization, excluding the last 15 min to avoid including the signal from contaminants/noise. This ‘MS2 signal’ of each file, akin to a total protein load stain on a Western blot gel, was used to adjust each transition intensity of each peptide in the corresponding file. Normalized transition-level data were then processed using mapDIA software to remove noisy/interference transitions from the peptide peak groups, calculate peptide and protein level intensities, and perform pairwise comparisons between groups.

The following pairwise comparisons were made: NOD vs. CD1 for each time point (weeks 10, 12, and 14); NOD-BDC2.5 vs. NOD-SCID ctrl; NOD resistant vs. NOD mice with diabetes. The mapDIA tool generated a q-value to indicate an FDR rather than a simple p-value. We assumed that protein expression differs significantly between two groups when the log_2_(fold-change) was >0.6 (i.e., ∼1.5 fold-change) and q-value/FDR was <0.01.

### Immunofluorescence staining and quantification

Immunofluorescence (IF) was performed to validate key findings from the MS analysis. Briefly, formalin-fixed paraffin-embedded (FFPE) pancreatic tissues from an independent cohort of pre-diabetic age-matched NOD and CD1 mice, obtained at 7, 9, 11, and 13 weeks of age, were sectioned at a thickness of 5 μm and deparaffinized. The sections were hydrated twice with fresh Xylene for 5 minutes and a series of decreasing ethanol concentration (100 to 70%). Antigen retrieval was performed using citrate buffer and stained using antibodies against PDIA1 (Cell Signaling, Cat# 3501S), PRDX3 (Abcam, Cat# ab73349), 14-3-3B/YWHAB (Sigma, Cat# HPA011212), insulin (Dako, Cat# IR002), and glucagon (Abcam, Cat# ab10988). Secondary antibodies included Alexa 488-labeled goat anti-guinea pig, Alexa 568-labeled donkey anti-rabbit, and Alexa 647-labeled donkey anti-mouse antibodies. Nuclei were stained with DAPI. Images were acquired using LSM800 confocal microscope (Zeiss, Germany) and quantified using image-J software (NIH) as described previously (69). From each animal, 3-7 islets were randomly selected for imaging. Normalized total islet cell fluorescence intensity was calculated independently by two individuals working in a blinded fashion.

### Collection of human serum samples

Fasting serum was obtained from children with recent-onset T1D and age- and sex-matched non-diabetic healthy controls under protocols approved by the Indiana University Institutional Review Board. Written informed consent or parental consent and child assent were obtained from all participants (72). Children with T1D had been newly diagnosed within 48 hours of blood collection and were hospitalized at the Riley Hospital for Children. Control pediatric subjects were ambulatory, did not take any chronic prescription medications, and were free of any chronic or acute illness within two weeks preceding sampling.

### Measurement of serum PDIA1

#### Assay development

To measure circulating levels of PDIA1, we developed a high-sensitivity electrochemiluminescence assay using the Meso Scale Discovery (MSD) ELISA conversion kit (Cat# K15A01-1), according to the manufacturer’s instructions. Briefly, five anti-P4HB/PDIA1 antibodies were purchased from multiple vendors (Table 2) and screened for their ability to bind human recombinant PDIA1 protein (rPDIA1). The day before the experiment, single spot standard plates of the conversion kit were washed three times with 150 μL of PBS and incubated overnight with 30 μL of each antibody in PBS at 4°C (73, 74). The following day, the antibodies were washed with 0.05% PBS-Tween 20 (PBS-T) and blocked with 1% of blocking buffer A (Cat# R93BA-1) for 1 hr in an orbital shaker at 700rpm. A 4-fold serial dilution of rPDIA1 was prepared with a starting concentration of 2500 ng/mL, which was added to the plates and incubated in an orbital shaker for 1 hr at RT. Then, the plates were washed three times with 0.05% PBS-T and incubated with a PDIA1 detection antibody generated from different species (for example, mouse capture antibody was used with rabbit detection antibody) to prevent cross-reactivity and in an orbital shaker for 1 hr. Subsequently, the plates were washed three times with 0.05% PBS-T and incubated with species-specific Sulfo-Tag for 1 hr in an orbital shaker. Next, the plates were washed three times with 0.05% PBS-T, 150 μL of 1X read-buffer (Cat # R92TC-2) was added to each well, and the signal was detected immediately using a MESO QuickPlex SQ 120 plate reader (MSD). Data were analyzed using Discovery Workbench software version 4.0.

**Table 1.**

|  | Age (years) | Gender (M/F) | BMI% | Diabetes (Yes/No) |
| --- | --- | --- | --- | --- |
| Healthy Controls | 12.1 ± 4.20 yrs | 7M/6F | 66.3 ± 24.33 | No |
| New Onset T1D | 11.57 ± 4.05 yrs | 6M/4F | 55.42 ± 28.37 | Yes |

**Table 2.**

| **Company** | **Catalogue Number** | **Antibody/recombinant protein** |
| --- | --- | --- |
| Sigma-Aldrich | HPA018884-100UL | Antibody |
| Thermo Scientific | Catalog # MA3-019 | Antibody |
| LSBio | LS-F72880-500 | P4HB ELISA Development Kit |
| Cell Signaling | Cat # 350 | Antibody |
| ProsPec | Cat#ENZ-252 | Recombinant protein |
| LSBio | Cat#LS-G22049 | Recombinant protein |

#### Validation in serum and plasma samples

Thirty μL of 2-fold diluted mouse plasma samples or thirty microliters of 4-fold diluted human serum samples were assayed, while 4-fold serially diluted rPDIA1 protein with a starting concentration of 2500 ng/mL was used to generate a standard curve. To quantitate circulating levels of PDIA1 in human serum and mouse plasma samples, standard one spot MSD plates were incubated with 5 μg/mL of capture antibody (Cat# HPA018884; Sigma) overnight at 4°C, and the same procedures described above under “*Assay development”* were followed. Following sample incubation, plates were washed as described above and incubated with mouse PDIA1 detection antibody (Cat# MA3-019; Thermo Fisher Scientific) for 1 hr in an orbital shaker at (RT). The plates were then washed and incubated with an MSD mouse Sulfo-Tag for 1 hr at RT in a shaker. Finally, the plates were read using 150 μL of read-buffer in a Quick Plex SQ 120 plate reader (MSD), and the data were analyzed as described above.

### Statistical analysis

Statistical analysis of the proteomics data is detailed above. Other experimental data were analyzed using GraphPad Prism version 9. Statistical significance of the difference between two groups was determined using the Student’s t-test and between more than two groups by one-way ANOVA; p≤0.05 was considered significant. Data are presented as mean ± S.E.M or mean ± S.D. Temporal changes in islet proteomics were analyzed using principal component analysis (PCA). Differentially expressed proteins were identified using unsupervised hierarchical clustering analysis and visualized using heatmap (https://software.broadinstitute.org/morpheus/). The upregulated/downregulated pathways were recognized using pathway enrichment and the Ingenuity pathway analysis.

## Contributors

FS and CEM conceived and designed the study. FS and RNB performed experiments. KR and J.V. prepared and ran LC-MS/MS samples. DS, XL, HW, and PK performed computational analyses. FS and CEM interpreted the data and wrote the manuscript. All authors provided critical revisions and edits to the manuscript. All authors approved the final manuscript. CEM is the guarantor of this work.

## Data Sharing Statement

Data presented as part of this manuscript are available from the corresponding author upon request.

## Funding and Acknowledgements

This work was supported by NIH grants R01 DK093954 and DK127308(to CEM) and U01DK127786 and UC4 DK 104166 (to RGM and CEM), VA Merit Award I01BX001733 (to CEM), 2-SRA-2019-834-S-B, JDRF 2-SRA-2018-493-A-B (to CEM and RGM), and gifts from the Sigma Beta Sorority, the Ball Brothers Foundation, and the George and Frances Ball Foundation (to CEM). F.S. was supported by JDRF postdoctoral fellowship (3-PDF-20016-199-A-N). The authors acknowledge the support of the Islet and Physiology Core and the Translation Core of the Indiana Diabetes Research Center (P30DK097512). We thank Dr. Emily Anderson-Baucum and Dr. Jan Kajstura for their assistance in editing the text of this manuscript.

