## Supplemental Figures and Legends for "A discovery-based proteomics approach identifies protein disulfide isomerase (PDIA1) as a biomarker of β cell stress in type 1 diabetes"

**Supplementary Figure Legends**

***Supplementary Figure 1S**: **Total number proteins identified between NOD and CD1 mice***. Ven diagram showing total number of common proteins identified between age- and sex-matched NOD and CD1 mice.

***Supplementary Figure 2S**: **Upregulated Pathways in NOD mice.*** Pathway enrichment analysis showing significantly upregulated pathways at week 10, 12, and 14 and at the time of diabetes onset in NOD mice.

***Supplementary Figure 3S**: **Downregulated Pathways in NOD mice.*** Pathway enrichment analysis showing significantly downregulated pathways at week 10, 12, and 14 and at the time of diabetes onset in NOD mice.

### Supplementary Figure 1S

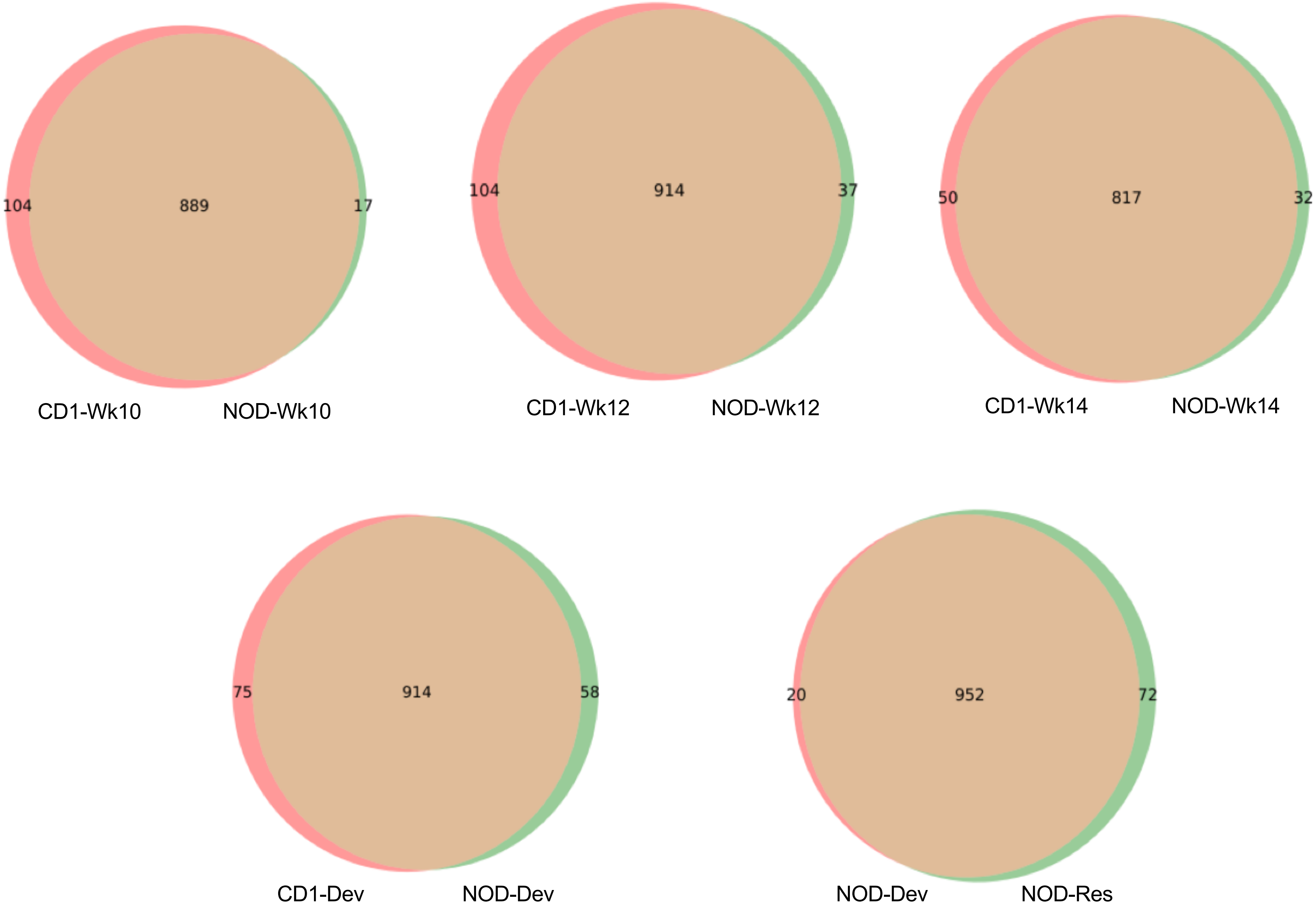

### Supplementary Figure 2S

#### Up-Regulated Pathways

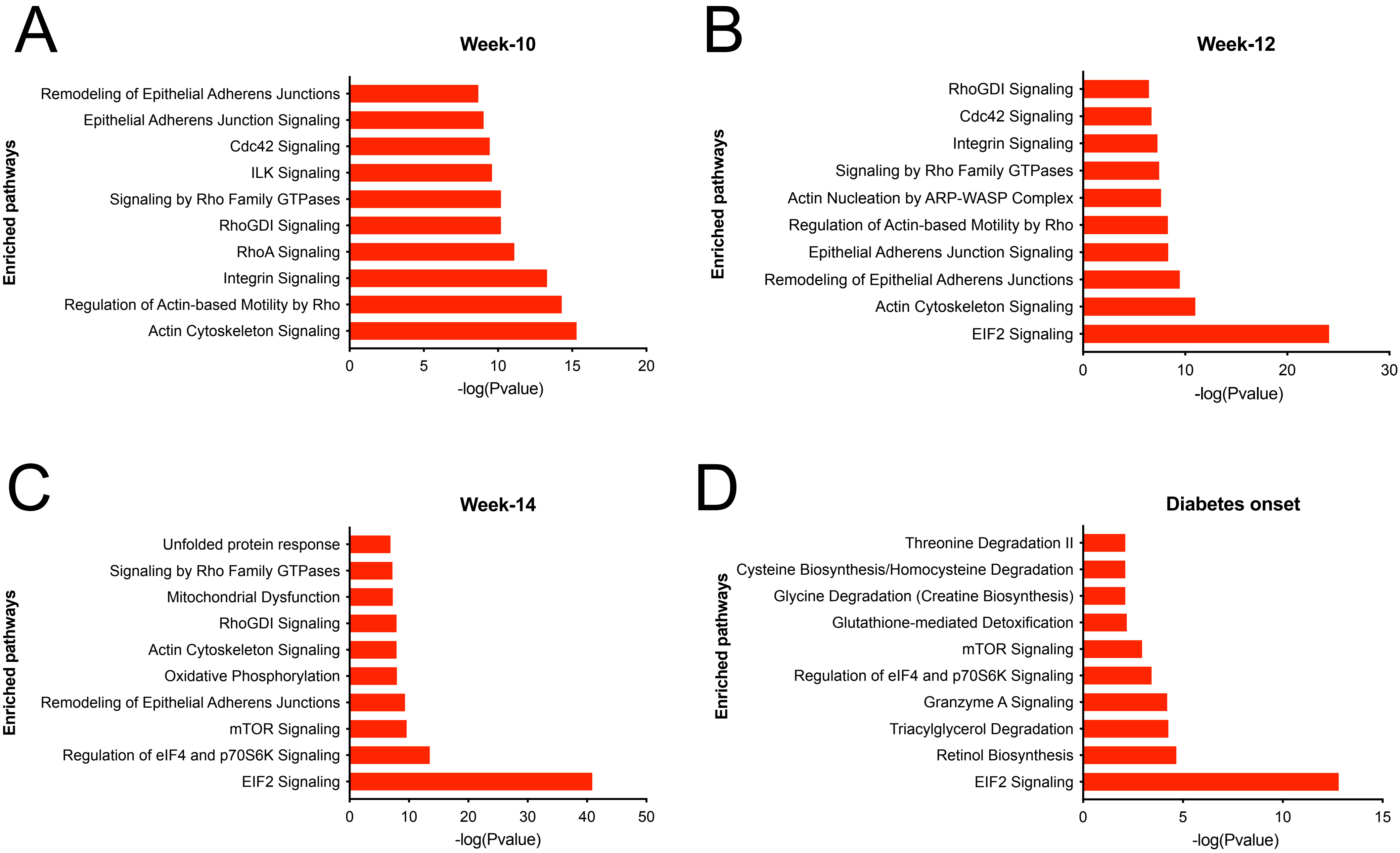

### Supplementary Figure 3S

#### Down-Regulated Pathways

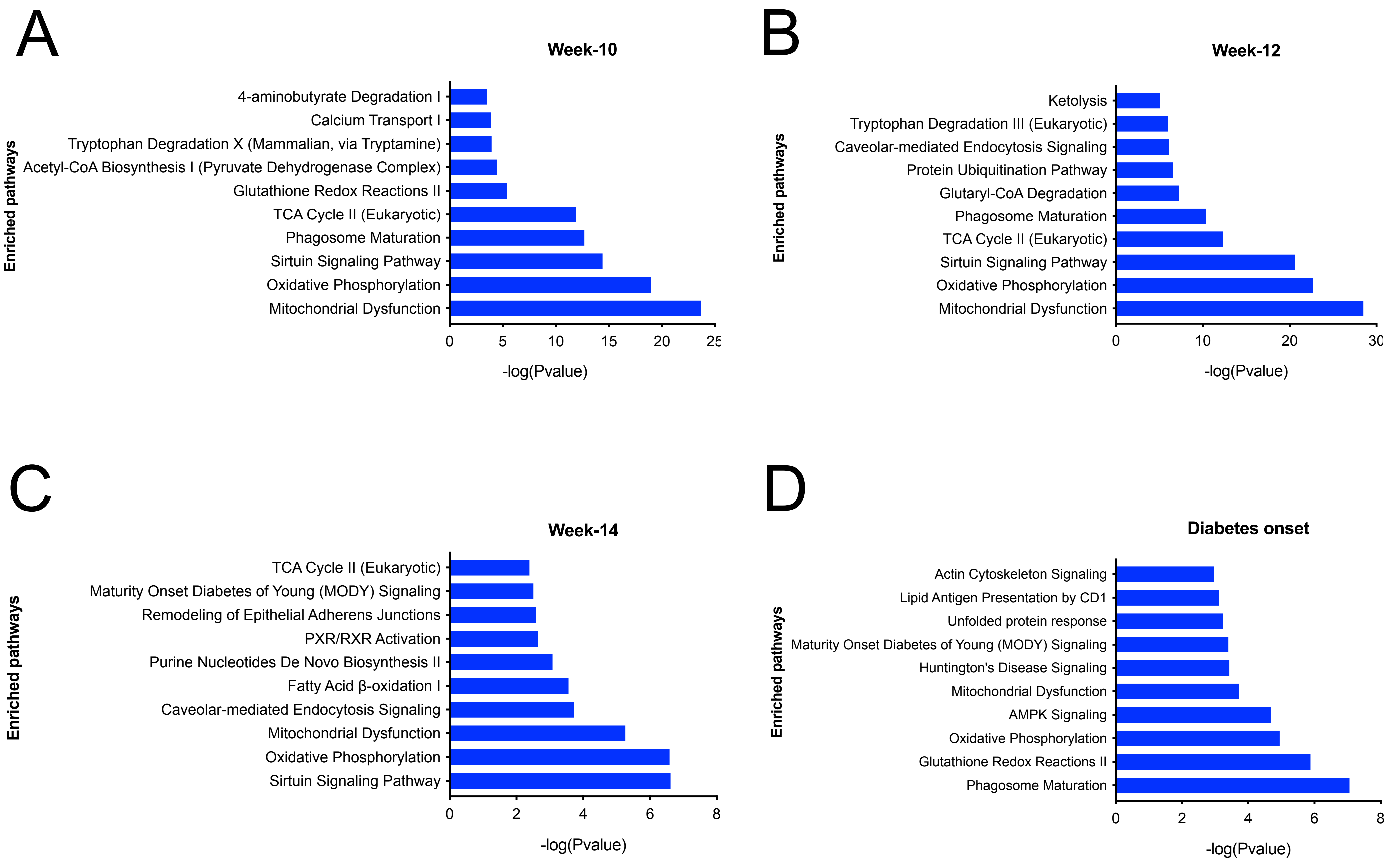
